# BuddySuite: Command-line toolkits for manipulating sequences, alignments, and phylogenetic trees

**DOI:** 10.1101/040675

**Authors:** Stephen R. Bond, Karl E. Keat, Sofia N. Barreira, Andreas D. Baxevanis

**Affiliations:** Computational and Statistical Genomics Branch, Division of Intramural Research, National Human Genome Research Institute, National Institutes of Health, 50 South Drive, Bethesda, MD, USA, 20892

**Keywords:** software, command line, sequence, alignment, phylogenetic tree, Python

## Abstract

The ability to manipulate sequence, alignment, and phylogenetic tree files has become an increasingly important skill in the life sciences, whether to generate summary information or to prepare data for further downstream analysis. The command line can be an extremely powerful environment for interacting with these resources, but only if the user has the appropriate general-purpose tools on hand. BuddySuite is a collection of four independent yet interrelated command-line toolkits that facilitate each step in the workflow of sequence discovery, curation, alignment, and phylogenetic reconstruction. Most common sequence, alignment, and tree file formats are automatically detected and parsed, and over 100 tools have been implemented for manipulating these data. The project has been engineered to easily accommodate the addition of new tools, it is written in the popular programming language Python, and is hosted on the Python Package Index and GitHub to maximize accessibility. Documentation for each BuddySuite tool, including usage examples, is available at http://tiny.cc/buddysuite_wiki. All software is open source and freely available through http://research.nhgri.nih.gov/software/BuddySuite.

## Introduction

Manipulation of biological sequence data is now a routine task within the life sciences, performed not just by bioinformaticians but also by ‘bench biologists’ who are becoming increasingly savvy in applying computational methods to their own work. While there are excellent graphical platforms for organizing, visualizing, and manipulating sequence data, it can be advantageous to interact with text files directly from the command line. Common tasks may include searching for specific records in a file, extracting subsequences, converting between formats, identifying motifs, or stripping poorly aligned regions from a multiple sequence alignment. While all of these can be accomplished with standard UNIX commands or existing open source software, combining tasks into a pipeline may require a series of intermediate files, custom command-line operations or scripts, and moving data between standalone tools or online services. As an alternative we present BuddySuite, a comprehensive set of general-purpose command-line tools for manipulating sequence, alignment, and phylogenetic tree data that can be joined into reproducible workflows using a simple unified syntax.

The European Molecular Biology Open Software Suite (EMBOSS) (Rice *et al.,* 2000) and Biopieces (www.biopieces.org) are the most comprehensive general-purpose open-source bioinformatics toolkits currently available for use at the command line. While both are excellent software packages, BuddySuite includes a number of new features we believe will benefit biologists, particularly those who are just starting to familiarize themselves with bioinformatic techniques. A notable change is our move away from the ‘one program per function' paradigm employed by EMBOSS and Biopieces; both of these packages contain over 200 separate programs, thus occupying a considerable namespace on a user's system. For comparison, the 105 function implemented in BuddySuite version 1.2+ are all contained within just four modules - SeqBuddy, AlignBuddy, PhyloBuddy, and DatabaseBuddy - with each module being responsible for a specific data type (i.e., sequences, alignments, phylogenetic trees, or online database queries, respectively). The first three modules rely on flags to differentiate among individual tools, providing a unified and intuitive interface. DatabaseBuddy, on the other hand, runs primarily as a ‘live shell’ where users interactively search and download sequence data stored in the NCBI, UniProt, and Ensembl public databases. In addition, file format detection is fully automated; any number of sequence, alignment, or phylogenetic tree files can be passed into their respective BuddySuite program, in any combination of supported formats, and the records will be parsed seamlessly (see Table 1 for a list of supported formats). This is particularly useful when using AlignBuddy or PhyloBuddy to call third-party alignment or phylogenetic inference programs, as any idiosyncratic format conversions are handled without the need of additional input from the user. All output is printed directly to the terminal window by default and each module adheres to the UNIX convention of accepting piped data, allowing individual tools to be daisy-chained into more complex workflows (illustrated further in the *Use-case examples* section below)

**Table 1.** File format support provided by each BuddySuite module for reading (R) and writing (W).

| Format | SeqBuddy | AlignBuddy | PhyloBuddy |
| --- | --- | --- | --- |
| Clustal | R & W<sup>†</sup> | R & W | None |
| EMBL<sup>‡</sup> | R & W | R<sup>†</sup> & W | None |
| FASTA | R & W | R<sup>†</sup> & W | None |
| GenBank<sup>‡</sup> | R & W | R<sup>†</sup> & W | None |
| Nexus | R & W<sup>†</sup> | R & W | R & W |
| Newick | None | None | R & W |
| NeXML | None | None | R & W |
| PHYLIP (interleaved) | R & W<sup>†</sup> | R & W | None |
| PHYLIP (sequential) | R & W<sup>†</sup> | R & W | None |
| SeqXML | R & W | None | None |
| Stockholm | R & W<sup>†</sup> | R & W | None |
| SWISS-PROT<sup>‡</sup> | R only | None | None |

<sup>†</sup>All sequences must be the same length<sup>‡</sup>Supports rich sequence annotation

One of the greatest advantages BuddySuite has over other programs is its handling of sequence feature annotation. Rich flatfile specifications such as GenBank and EMBL support annotation, but this information is generally discarded by EMBOSS, and Biopieces is unable to create new records in these formats itself. BuddySuite modules recognize features in sequence files during processing and will update those annotations if a sequence is modified. For example, when a GenBank cDNA file is translated to protein, the relative position of each feature is scaled by one-third to account for the conversion of codons to amino acids. Similarly, if those protein sequences are passed into a supported multiple sequence alignment program, such as MAFFT (Katoh and Standley, 2013), the feature positions will be adjusted to account for gaps.

## Use-case examples

The use cases highlighted in this section are intended to illustrate common features of the BuddySuite modules. For extended documentation, please refer to the public wiki (http://tiny.cc/buddysuite_wiki), where each tool is described in greater detail. Furthermore, the module names in the examples below have been shortened to sb for SeqBuddy, alb for AlignBuddy, and pb for PhyloBuddy.

BuddySuite modules are executed from the command line using the following generalized syntax:

~~~
$: module file(s) <cmd> <args> <modifiers>
~~~

Any number of files may be passed into the module, followed by a flag to specify which command to run and any additional arguments that command requires. As a specific example, the following would accept two sequence files, one in FASTA format and the other in GenBank format, then delete sequences larger than 300 residues:

~~~
$: sb seqsl.gb seqs2.fa --delete_large 300
~~~

Whenever multiple file specifications are combined like this, the rightmost file will determine the final output format (in this case, FASTA), although this behaviour may be overridden with the ‘--output’ modifier. The command below would yield records in GenBank format:

~~~
$: sb seqs1.gb seqs2.fa —delete_large 300
 --output genbank
~~~

Modifiers like ‘--output’ are used sparingly in BuddySuite and only when their effects are intuitively applicable across most tools (e.g., ‘--quiet’ execution or to rewrite files ‘--in_place’).

Complex workflows can also be built from the BuddySuite modules using the pipe character:

~~~
$: sb cDNA_seqs.fa —transmembrane_domains |
 sb --pull_records "TMD4" |
 sb --translate |
 alb --generate_alignment mafft |
 alb --extract_feature_sequences "TMD2:TMD4" |
 pb --generate_tree raxmlHPC-SSE3 |
 pb --root
~~~

In the example above, SeqBuddy processes a set of homologous cDNAs and transmits them to the TopCons server (Tsirigos *et al.,* 2015) to predict transmembrane domains (TMD). Once the results have been automatically retrieved from the server, the sequences are annotated and those that include ‘TMD4’ are retained for further analysis (note that SeqBuddy recasts the format to GenBank when applying new sequence features). The cDNAs are translated to protein, a multiple sequence alignment is generated, the section of the alignment spanning the second through forth TMD is extracted, and then phylogenetic inference software is called via PhyloBuddy to generate a tree. Finally, the tree is rooted at its midpoint.

To piece this workflow together without BuddySuite, a user may identify TMDs with a command-line version of software like TMHMM (Krogh *et al.*, 2001) or access the TOPCONS server directly from a web browser, and then manually inspect the output files or parse them with awk to create a list of appropriate sequence IDs. After pulling those sequences from the original file with seqret and translating them with transeq (which are both part of the EMBOSS suite), they would then need to be saved as a new FASTA file in order to be passed into MAFFT to generate the multiple sequence alignment. Extracting the alignment columns covering TMD2 through TMD4 would require further manual inspection or a custom script to match the location of each TMD from the TMHMM or TOPCONS results to the alignment. Once identified, seqret could be called again to pull the correct regions from the alignment and create a new file. Finally, RAxML (Stamatakis, 2006) could be used to infer a phylogenetic tree and root it. While the above series of steps is certainly valid and will accomplish the end goal of the user, BuddySuite offers a far simpler solution involving fewer pieces of software, fewer intermediary files, less manual intervention, and a consistent syntax.

## Installation

The BuddySuite libraries have been written in Python 3 for use on all major operating systems (Windows 7+, Mac OSX, and Linux). Stable release versions can be installed directly from the Python Package Index (PyPI) using the popular package manager ‘pip’, and the most recent development version is continually available from GitHub. Dependencies have been limited to packages available through PyPI to simplify installation, although a number of optional third-party programs can also be accessed through BuddySuite; these include BLAST (Camacho *et al.,* 2009) for comparing sequences, multiple sequence alignment packages like MAFFT, and phylogenetic inference packages like RAxML. Installation of these programs (itemized in Table 2) is discretionary because they are not necessary for the general operation of the BuddySuite modules. Users are also encouraged to run the optional BuddySuite setup script after installation to configure their preferences:

~~~
$: pip install buddysuite
$: buddysuite setup
~~~

**Table 2.** List of optional third party software that BuddySuite programs can interact with.

|  | Program | Reference |
| --- | --- | --- |
| SeqBuddy | BLAST | (Camacho et al., 2009) |
| AlignBuddy | ClustalΩ | (Sievers et al., 2011) |
|  | ClustalW2 | (Larkin et al., 2007) |
|  | MAFFT | (Katoh and Standley, 2013) |
|  | MUSCLE | (Edgar, 2004) |
|  | PAGAN | (Löytynoja et al., 2012) |
|  | PRANK | (Löytynoja and Goldman, 2005) |
| PhyloBuddy | FastTree | (Price et al., 2010) |
|  | RAxML | (Stamatakis, 2006) |
|  | PhyML | (Guindon et al., 2010) |

BuddySuite performs all necessary format conversion to call any of these tools and, where appropriate, returns the result in the same format as the input. This is particularly useful when creating multiple sequence alignments from annotated sequences in GenBank or EMBL format.

## Developers

Looking forward, the modular nature of BuddySuite makes it particularly well-suited to open-ended development. New tools are easily added to each existing module and new modules may eventually extend the suite to new data types. While we will continue our own efforts to support and expand BuddySuite, we also strive to attract input and contributions from the broader biomedical community. To minimize barriers against community-driven development, the project is maintained on GitHub, has comprehensive unit test coverage of over 95%, includes extensive and accessible documentation, and makes every effort to conform with open-source best practices (Leprevost *et al.,* 2014; Seemann, 2013).

Instead of relying exclusively on active input from users, we have implemented an optional passive data collection system to monitor usage and report crashes. If activated during setup, this software improvement program will periodically transmit anonymized data to the core developers. In the event of a crash, it will analyze the traceback and compare it against an online database of resolved issues before immediately informing the user if the problem is patched by a newer release.

The internal functionality of BuddySuite is also easily accessible by developers wishing to write third-party Python programs. Each module has a core ‘Buddy’ class that automatically processes a variety of input types, including plain text, file paths, file handles, and lists of record objects. They also perform all necessary file format processing and expose methods for managing and writing the sequence or tree records. The functions in each library accept these Buddy objects as input and generally return them as output, thus providing a standardized application programing interface that facilitates interoperability among functions. Once installed, the BuddySuite libraries can be imported using conventional Python syntax.

## Conclusions

BuddySuite has been designed from the ground up as an intuitive, extensible, and unified platform for routine command-line tasks performed on sequence, alignment, and phylogenetic tree files. This is the first time such a large suite of general-purpose bioinformatics utilities have been implemented purely in Python and packaged together under a flag-driven paradigm. Well-designed and actively supported open-source tools will be invaluable over the coming years as an increasing number of biologists without formal bioinformatics training turn to the command line to analyze their data. By combining active community feedback with the passive data-collection features built into this project, we look forward to tailoring future development to the needs of our users.

## Acknowledgments

This research was supported by the Intramural Research Program of the National Human Genome Research Institute, National Institutes of Health. We would also like to thank Drs. Maxence LeVasseur and Tyra Wolfsberg for their thoughtful feedback on this manuscript and the community members who contributed code to the project, big or small. It takes a village.

